# The risk of sustained sexual transmission of Zika is underestimated

**DOI:** 10.1101/090324

**Authors:** Antoine Allard, Benjamin M. Althouse, Laurent Hébert-Dufresne, Samuel V. Scarpino

**Affiliations:** Departament de Física de la Matèria Condensada, Universitat de Barcelona, Martí i Franquès 1, E-08028 Barcelona, Spain; Universitat de Barcelona Institute of Complex Systems (UBICS), Universitat de Barcelona, Barcelona, Spain; Institute for Disease Modeling, Bellevue, WA, 98005, USA; University of Washington, Seattle, WA, 98195, USA; New Mexico State University, Las Cruces, NM, 88003, USA; Santa Fe Institute, Santa Fe, NM, 87501, USA; Department of Mathematics and Statistics and Complex Systems Center, University of Vermont, Burlington, VT, USA

## Abstract

Pathogens can follow more than one transmission route during outbreaks – from needle sharing plus sexual transmission of HIV to small droplet aerosol plus fomite transmission of influenza. Thus, controlling an infectious disease outbreak often requires characterizing the risk associated with multiple mechanisms of transmission. For example, during the Ebola virus outbreak in West Africa, weighing the relative importance of funeral versus health care worker transmission was essential to stopping disease spread [1]. Strategic policy decisions regarding interventions must rely on accurately characterizing risks associated with multiple transmission routes. The ongoing Zika virus outbreak challenges our conventional methodologies for translating case-counts into route-specific transmission risk. Critically, most approaches will fail to accurately estimate the risk of seeing sustained sexual transmission of a pathogen that is primarily vectored by a mosquito – such as the case with the risk of sustained sexual transmission of Zika virus [2, 3].

In conventional models of disease transmission, if the basic reproductive number of a pathogen, *R*_0_ (the average number of secondary infections resulting from a single infected individual in an entirely susceptible population), is less than one, then a large outbreak is unlikely. Therefore, public health officials are interested in both estimating *R*_0_ and evaluating which interventions are most likely to reduce it. Having *R*_0_ *≥* 1 is a necessary condition to obtain a large epidemic, and it relies on the assumption that the infection follows a branching process. With *R*_0_ *≥* 1, each infected individual will on average give rise to at least one additional case and the outbreak will be self-sustaining in a large population; while the branching process dies out if *R*_0_ *<* 1 as individuals give rise to less than one additional case on average. In the unique case of sexual transmission of Zika, this underlying assumption is broken in three ways: (1) individuals infected by the vector are not necessarily part of the branching process over the network of sexual contacts; (2) all infections are not equivalent since there is a strong asymmetry in sexual transmission as men are infectious for much longer period than women [4]; and (3) symptoms, and thus probability of detection, are also heterogeneous [5].

First, while it is tempting to evaluate the sexual *R*_0_ of Zika by averaging the number of sexual transmissions caused by all known infected individuals, it is not the correct approach. To estimate the *R*_0_ for sexual transmission correctly, we need to evaluate the number of secondary infections caused by individuals who were infected sexually. The idea is that individuals infected by a mosquito were randomly drawn from the total population (and thus have a number of sexual partners drawn from some distribution {*p_k_*}), whereas individuals infected sexually are conditioned on having at least one contact and are in fact 10 times more likely to have degree 10 than degree 1 (their number of sexual partners is thus drawn from a distribution proportional to {*kp_k_*}). This distinction is the source of a classic network theory result: your friends have more friends than you do [6].

For a sexually transmitted infection on a contact network, this means that the key quantity is not the expected number of contacts an individual has, i.e the degree of a node, but is instead the expected *excess* number of contacts: the number of edges other than the one through which they were infected [7]. In contrast, for individuals who are infected by a mosquito we must use the expected number of contacts, because the infection was transmitted to them by the vector and not by one of their sexual/social contacts (see Fig. 1).

**Figure 1.**
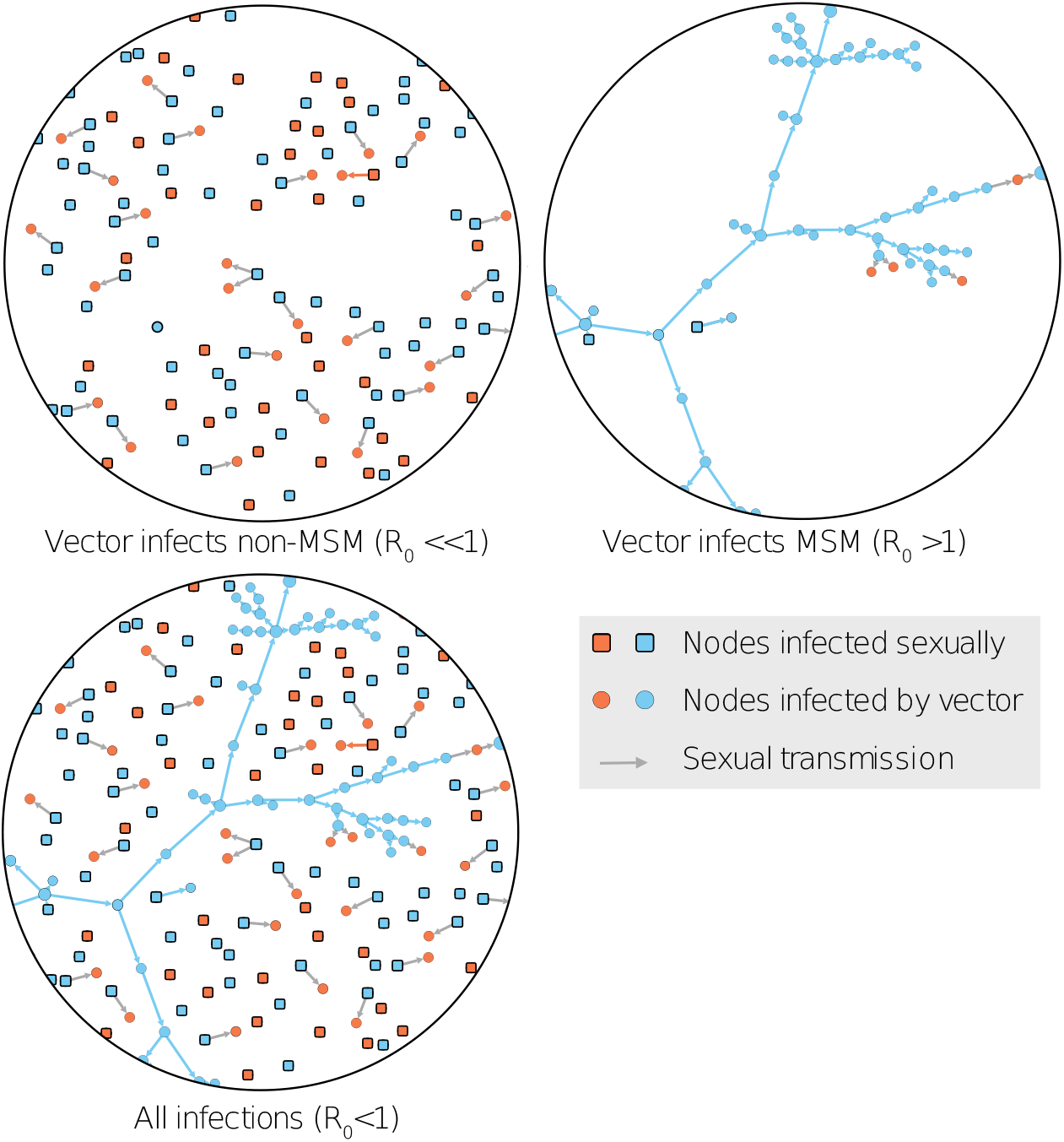
Orange nodes are female, blue nodes are male. Most individuals will be infected by the vector and typically infect 0 or 1 sexual partners. However, within the men who have sex with men community, the probability of transmission is such that an epidemic is possible (with some spillover transmission to heterosexual individuals through bisexual individuals).

If probability of sexual transmission is low, then the average number of sexual transmissions an individual causes is close to what we would expect given the average number of sexual partners. However, if most of the infections in the population are due to sexual transmissions, then the average number of sexual transmissions an individual causes is close to what we would expect given the average excess number of sexual partners. We expect this second average to be larger. As such, measurements of the typical number of sexual transmissions caused by an individual in a population with many vector-caused infections will underestimate the typical number of sexual transmissions caused by an individual in a population with few or no vector-caused infections (see the left panel of Fig. 2).

**Figure 2.**
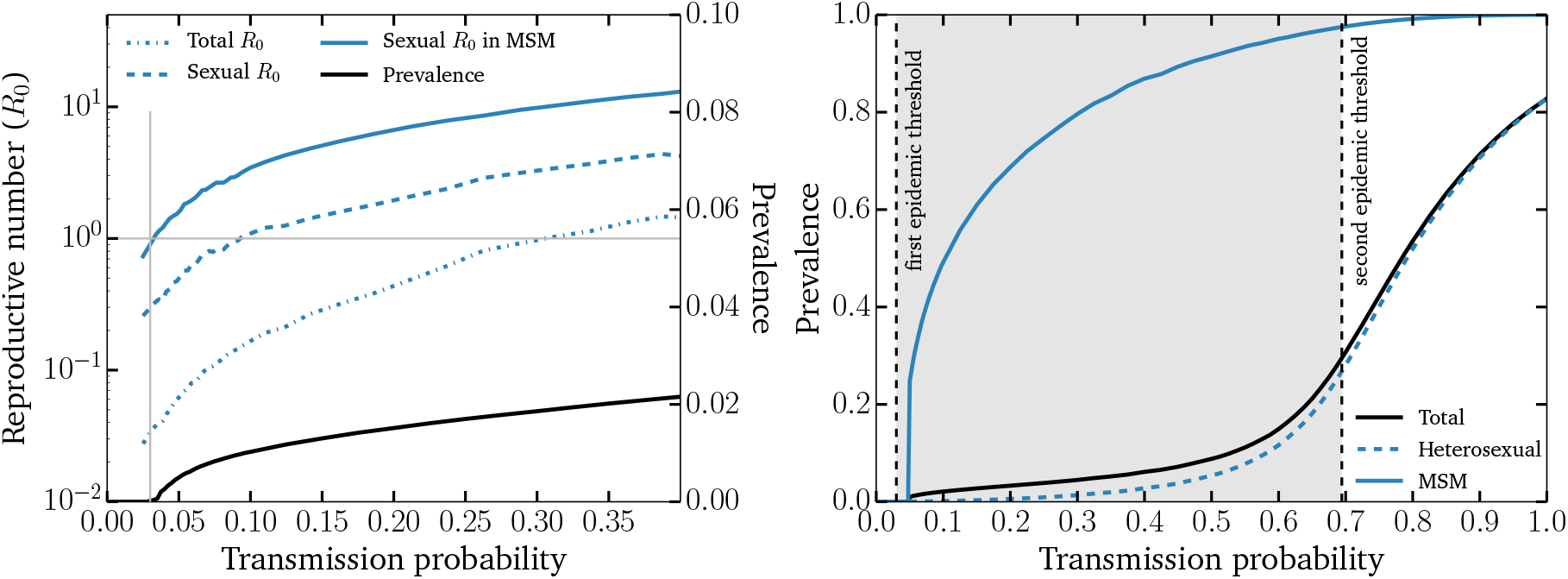
We simulate Zika transmission on a random network of sexual contacts where the distribution of contacts are drawn from Ref. [8] (for homosexual contacts) and Ref. [9] (for heterosexual contacts). A population of 10^6^ individuals is assumed to be equally parted between men and women, of which 5% are homosexuals, 3% are bisexuals and 92% are heterosexuals. We assume very few individuals are infected by the mosquito vector, but they can cause outbreaks through sexual transmission where men transmit with probability *T* and women with probability *εT*. (left) Using *ε* = 1/10 and the full population, we show three estimates of the basic reproductive number *R*_0_ caused by sexual transmission: the total *R*_0_ is the average number of secondary infections for all infected individuals, the sexual *R*_0_ is the average number of secondary infections for individuals infected sexually, and the same sexual *R*_0_ only for MSM infected sexually. Using all data points to estimate *T_c_* through *R*_0_ *=* 1 leads to overestimation of *T_c_* by a factor of 10 (0.30 versus 0.03 as the true sexual *R*_0_ in the MSM population). (right) We show the total prevalence of Zika (and the prevalence in two sub-populations) as a function of T using *ε* = 1/2 and only individuals with more than one sexual partner (as a rough approximation of the sexually active population). Also shown are the threshold values for a self-sustained epidemic *within* the MSM sub-population (with subcritical spillovers in the rest of the population, shaded area), and within the whole population (i.e., supercritical outbreaks exist *outside* of the MSM sub-population).

Second, even ignoring complications due to the vector, the sexual transmission of Zika is itself complicated: males and females are not equally likely to transmit infection due to differences in duration of Zika virus in semen and vaginal fluids. In fact, females are significantly less likely to transmit Zika to their sexual partner than males [4, 10]. Because of the asymmetry in infectious period between males (> 180 days [4]) and females (< 20 days [10]), it is much easier for Zika to invade the men who have sex with men (MSM) community. Consider two infectious males, one in the MSM community and one not in the MSM community: Zika only needs to be transmitted to at least one other individual on average to permit an epidemic in the MSM community, whereas in the non-MSM community, Zika needs to infect enough females such that they will together infect at least one other male. The stronger the asymmetry in infectious period, the more different these conditions become. There are thus two different epidemic thresholds [11]: one for a large-scale epidemic in the MSM community (around which small *subcritical* outbreaks will spillover in the rest of the population); and a higher transmission threshold for an epidemic in the entire population (see the right panel of Fig. 2).

Combining the effects of the multiple transmission pathways and the asymmetric transmission explains the results of shown in the left panel of Fig. 2. For Zika, the real sexual *R*_0_ must be estimated by considering only MSM (because of its asymmetry in transmission) and only those infected sexually (because of its multiple transmission pathways). The mathematical treatment of the asymetric transmission is detailed in Ref. [11] and that of the multiple transmission pathways in Ref. [12].

Third, an insidious aspect of the double epidemic threshold of Fig. 2 is that one of the most likely outcomes – an epidemic mostly contained within the MSM community – is also the hardest outcome to detect. Zika virus infections in adults are largely asymptomatic [5] and, therefore, most testing occurs in the roughly 20% of cases that are symptomatic or in individuals seeking to have children [13]. The vast majority of these individuals will be outside of the MSM community [13]. As a result, the group most likely to see sustained sexual transmission is the group least likely to be tested. Given that most infections are asymptomatic in adults [5, 13], this lack of awareness seems less critical; however, it will clearly lead to biased estimates in the potential for sustained sexual transmission of Zika. Sustained sexual transmissions in the MSM community will lead to sporadic sub-critical but potentially dramatic spillover in the heterosexual community. Hence, given our evolving understanding of the clinical outcomes for Zika infection in adults, it may be prudent from an individual perspective, as well as a population-level perspective, to increase testing for all adults.

Public health decision- and policy-makers rely on accurate characterizations of transmission risk to decide on interventions and allocate strategic resources. For pathogens, like Zika, which are both vectored by an insect and transmitted sexually, conventional approaches will underestimate the risk associated with sexual transmission. Underestimating the risk of sexual transmission will lead both to biased intervention efforts and to an underestimate of the potential for disease persistence. Additionally, given the differential rates of Zika testing in MSM versus non-MSM communities, the data on sexual transmission are almost certainly biased. Here we advocate for the decision-making and disease modeling communities to better embrace contact network methods when characterizing transmission risk and for increased testing in both MSM and non-MSM communities. The resulting estimates of transmission risk and key epidemiological measures, e.g. *R*_0_ and the likelihood of disease persistence, will be more accurate.

## Acknowledgments

The authors thank Joel C. Miller for useful comments on the manuscript. A.A. acknowledges support from the Fonds de recherche du Québec – Nature et technologies. S.V.S acknowledges funding from the University of Vermont. L.H.-D. acknowledges the Santa Fe Institute and the James S. McDonnell Foundation Postdoctoral Fellowship. B.M.A. and L.H.-D. thank Bill and Melinda Gates for their support of this work and their sponsorship through the Global Good Fund.

## Author contributions

A.A. and L.H.D performed the calculations and simulations; all authors conceived of the project, interpreted the results, and produced the final manuscript. Authors are listed alphabetically.

